# Enhanced transmissibility of XBB.1.5 is contributed by both strong ACE2 binding and antibody evasion

**DOI:** 10.1101/2023.01.03.522427

**Authors:** Can Yue, Weiliang Song, Lei Wang, Fanchong Jian, Xiaosu Chen, Fei Gao, Zhongyang Shen, Youchun Wang, Xiangxi Wang, Yunlong Cao

**Author notes:** Correspondence: Xiangxi Wang, Yunlong Cao. These authors contributed equally (Can Yue, Weiliang Song).

## Abstract

SARS-CoV-2 recombinant subvariant XBB.1.5 is growing rapidly in the United States, carrying an additional Ser486Pro substitution compared to XBB.1 and outcompeting BQ.1.1 and other XBB sublineages. The underlying mechanism for such high transmissibility remains unclear. Here we show that XBB.1.5 exhibits a substantially higher hACE2-binding affinity compared to BQ.1.1 and XBB/XBB.1. Convalescent plasma samples from BA.1, BA.5, and BF.7 breakthrough infection are significantly evaded by both XBB.1 and XBB.1.5, with XBB.1.5 displaying slightly weaker immune evasion capability than XBB.1. Evusheld and Bebtelovimab could not neutralize XBB.1/XBB.1.5, while Sotrovimab remains weakly reactive and notably, SA55 is still highly effective. The fact that XBB.1 and XBB.1.5 showed comparable antibody evasion but distinct transmissibility suggests enhanced receptor-binding affinity would indeed lead to higher growth advantages. The strong hACE2 binding of XBB.1.5 could also enable its tolerance of further immune escape mutations, which should be closely monitored.

## Main

SARS-CoV-2 subvariants BQ.1.1 and XBB.1 have been circulating globally with superior growth advantages over most Omicron mutants (Fig. S1A). However, XBB.1.5, a subvariant of the recombinant mutant XBB, has recently shown a substantial growth advantage over BQ.1.1 and XBB.1. Given its enhanced transmissibility, XBB.1.5 has rapidly become the dominant strain in the United States and is highly likely to cause the next global wave of COVID-19 (Fig. S1B) ^1^. XBB/XBB.1 is already demonstrated to be extremely evasive against the neutralization of plasma/serum from vaccinated or convalescent individuals and monoclonal antibodies (mAbs), even stronger than that of BQ.1.1 ^2-5^. Compared to XBB.1, XBB.1.5 carries a Ser486Pro mutation on the spike protein, a rare 2-nucleotide substitution compared to the ancestral strain (Fig. S1C). The mechanism behind the rapid transmission of XBB.1.5, especially the impact of Ser486Pro, requires immediate investigation.

Here, using vesicular stomatitis virus (VSV)-based pseudovirus neutralization assays, we evaluated the neutralization titers against XBB.1.5 of convalescent plasma from individuals who had received 3 doses of CoronaVac prior to BA.1, BA.5, or BF.7 breakthrough infection (BTI). A cohort of convalescents from BA.5 BTI who had received at least two doses of BNT162b2 or mRNA-1273 is also included in the analysis. Human ACE2 (hACE2)-binding affinity of XBB.1.5 receptor-binding domain (RBD) was also examined by surface plasmon resonance (SPR) assays, in comparison to that of XBB.1, BQ.1.1, and BA.2.75. Plasma samples associated with CoronaVac were collected on average 4 weeks after hospital discharge (Table S1). Plasma samples associated with the mRNA vaccine were collected within 2-3 weeks after hospital admission (Table S1). The lack of BQ.1.1 BTI convalescent individuals is a limitation of this study to estimate the scale of immune evasion of XBB.1.5 for this group.

Plasma samples from CoronaVac-vaccinated BA.1/BA.5/BF.7 BTI showed a substantial decrease in plasma 50% neutralization titer (NT50) against XBB.1 and XBB.1.5 compared to that against B.1 (D614G) variant (Fig. 1A). Specifically, plasma from post-CoronaVac BA.5 BTI showed a 44-fold decrease in NT50 against XBB.1 compared to that against B.1, while the decrease is 40-fold for XBB.1.5. For post-CoronaVac BF.7 BTI, the plasma NT50 against XBB.1 and XBB.1.5 was decreased 31 and 27-fold respectively compared to that against B.1. A similar trend was also observed in plasma from mRNA-vaccinated BA.5 BTI and post-CoronaVac BA.1 BTI. The above results indicate that Pro486 is also a strong neutralizing antibody evading mutation, and the humoral immune escape ability of XBB.1.5 is comparable to XBB.1, in fact slightly weaker.

**Figure 1.**
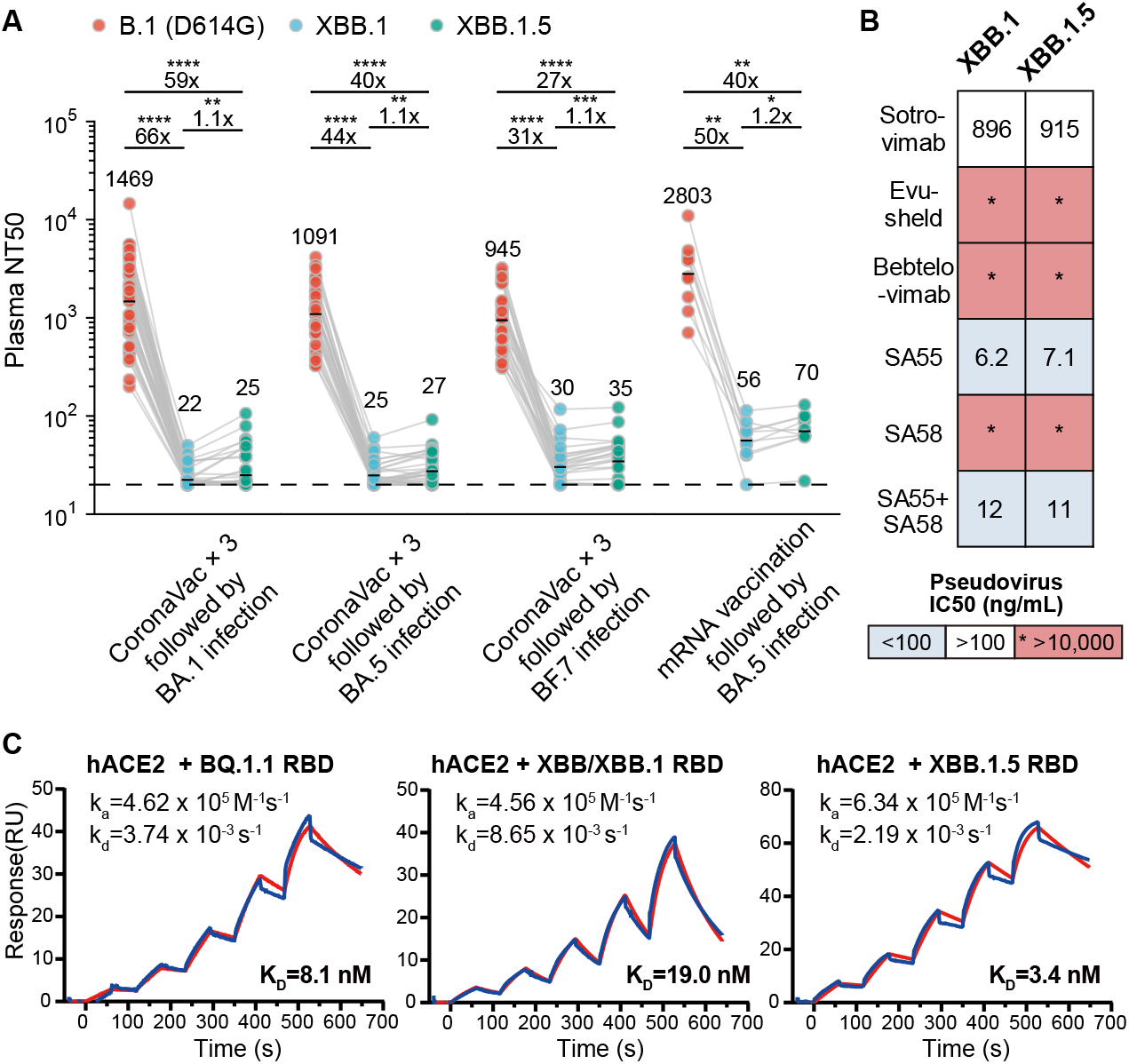
XBB.1.5 exhibits enhanced hACE2 binding with strong antibody evasion. (A) NT50 against SARS-CoV-2 B.1 (D614G), XBB.1, and XBB.1.5 pseudovirus using plasma from BA.1 (n=50), BA.5 (n=36), or BF.7 (n=30) BTI convalescents with 3 doses of CoronaVac in prior, and BA.5 BTI convalescents with 3 or 4 doses of vaccination in prior including at least two doses of mRNA vaccines (BNT162b2 or mRNA-1273) (n=10). P-values were calculated using two-tailed Wilcoxon signed rank tests. *p<0.05; **p<0.01; ***p<0.001; ****p<0.0001; NS, not significant. All neutralization assays were conducted in at least two independent experiments. (B) Pseudovirus 50% inhibition concentration (IC50) of therapeutic neutralizing antibodies. Blue, white, and red backgrounds indicate IC50 < 100ng/mL, > 100ng/mL, and > 10,000ng/mL (limit of detection, marked as an asterisk), respectively. All neutralization assays were conducted in at least two independent experiments. (C) SPR sensorgrams measuring the hACE2-binding affinity of SARS-CoV-2 BQ.1.1, XBB/XBB.1, and XBB.1.5 RBD. The fitted association rate constants (k_a_), dissociation rate constants (k_d_), and dissociation equilibrium constants (K_D_) are shown.

Compared to XBB.1, XBB.1.5 exhibited similar evasion against therapeutic mAbs (Fig. 1B). Evusheld and Bebtelovimab could not neutralize XBB.1.5 pseudovirus. S309 is still active but weak against XBB.1.5. SA58 is escaped, while SA55 remains highly effective against XBB.1.5 ^2,6^.

Previous deep mutational scanning studies have shown that Pro486 may enhance the affinity to human ACE2 (hACE2) compared to Ser486 ^7^. Indeed, the hACE2-binding affinity of XBB.1.5 RBD (with a dissociation constant K_D_ of 3.4 nM) was comparable to that of BA.2.75 (K_D_=1.8 nM) and much stronger than that of XBB.1 (K_D_=19 nM) and BQ.1.1 (K_D_=8.1 nM) (Fig. 1C and S2). These results suggest that the probable reason for the significant growth advantage of XBB.1.5 over XBB.1 is that it gained substantially higher ACE2 binding affinity through the Ser486Pro mutation, while retaining an extremely high immune evasion capability.

With stronger immune escape ability than BQ.1.1 but limited by weaker ACE2 binding affinity, XBB and XBB.1 have only prevailed in a few countries, such as Singapore and India, in the past few months, while BQ.1.1 has quickly become the global dominant strain. Given its enhanced hACE2-binding affinity but comparable antibody evasion, the prevalence of XBB.1.5 demonstrates that receptor-binding affinity will substantially affect the transmissibility, but the underlying mechanism still needs further investigation. Also, whether the increased receptor-binding affinity would cause a difference in pathogenicity compared to XBB is unclear and requires immediate research ^8^. Moreover, the strong affinity to hACE2 may allow XBB.1.5 to acquire additional immune-escape mutations, similar to the evolution trend of BA.2.75, when met with substantial immune pressure ^9^. Therefore, the circulation of XBB.1.5 needs to be closely monitored, and the development of effective neutralizing antibodies and vaccines against XBB.1.5 is urgently needed.

## Declaration of interest

Y.C. is a co-founder of Singlomics Biopharmaceuticals and inventor of provisional patents associated with SARS-CoV-2 neutralizing antibodies, including SA55 and SA58. All other authors declare no competing interests.

## Acknowledgments

We thank all volunteers for providing the blood samples. We thank all the scientists around the globe for performing SARS-CoV-2 sequencing and surveillance analysis. This project is financially supported by the Ministry of Science and Technology of China and Changping Laboratory (2021A0201; 2021D0102), and the National Natural Science Foundation of China (32222030).

## Supplementary Appendix

### Author Contributions

Y.C. and X.W. designed and supervised the study. W.S., F.J. and Y.C. wrote the manuscript. C.Y., L.W., and X.W. performed and analyzed the SPR experiments. W.S., F.J. Y.W., and Y.C. performed and analyzed the pseudovirus neutralization assay. X.C., F.G., and Z.S. recruited the COVID-19 convalescent individuals.

### Ethic Statement

This study was approved by the Ethics Committee of the Fourth Hospital of Inner Mongolia (Ethics committee archiving No. 202220), the Ethics Committee of Tianjin First Central Hospital (Ethics committee archiving No. 2022N045KY), and the Ethic Committee of Beijing Ditan Hospital, Capital Medical University (Ethics committee archiving No. LL-2021-024-02). Informed consent was obtained from all human research participants.

### Methods

#### Plasma isolation

Blood samples from individuals who had post-vaccination BA.1, BA.5, or BF.7 infection were collected under study protocols approved by the Ethics Committee of the Fourth Hospital of Inner Mongolia (Ethics committee archiving No. 202220), the Ethics Committee of Tianjin First Central Hospital (Ethics committee archiving No. 2022N045KY), and the Ethic Committee of Beijing Ditan Hospital, Capital Medical University (Ethics committee archiving No. LL-2021-024-02). Detailed information about the vaccination and infection status of each individual is available in Table S1. The infected strains of each group of convalescent patients were either confirmed by sequencing, or epidemiologically linked to other confirmed patients during a regional outbreak in China. Blood samples were diluted 1:1 with PBS+2% FBS (Gibco) and subjected to Ficoll (Cytiva) gradient centrifugation. Plasma was collected from the upper layer and stored at -80 °C until use.

#### Expression of monoclonal neutralizing antibodies

Heavy and light chain sequences of Evusheld (COV2-2196/COV2-2130, PDB: 7L7E) ^10^ and Bebtelovimab (LY-CoV1404, PDB: 7MMO) ^11^ were downloaded from PDB. Sequences of SA55 and SA58 have been reported previously ^6^. The antibodies were expressed as human IgG1 using Expi293F™ (ThermoFisher) cell lines in house, and purified with Protein-A magnetic beads (Genscript), as described previously ^2^.

#### Pseudovirus neutralization assay

The sequence of the S protein of SARS-CoV-2 ancestral strain (GenBank: MN908947) with specific mutations was codon-optimized for mammalian cells and inserted into the pcDNA3.1 vector. Site-directed mutagenesis PCR was performed as described previously ^12^. Detailed mutations carried by variants involved in this study are shown in Figure S1C in the Appendix. 293T cells (ATCC, CRL-3216) were transfected with pcDNA3.1-Spike using Lipofectamine 3000 (Invitrogen). The transfected cells were subsequently infected by G*ΔG-VSV (Kerafast) packaging expression cassettes for firefly luciferase instead of VSV-G in the VSV genome. Supernatants were discarded after 6-8h harvest and replaced with complete culture media. The cells were cultured for 1 day, and then the cell supernatants containing the spike-pseudotyped virus were harvested, filtered (0.45-μm pore size, Millipore), aliquoted, and stored at -80 °C. Viruses of multiple variants were diluted to the same number of copies before usage.

For neutralization assays, plasma or mAbs were serially diluted and incubated with the pseudovirus in 96-well plates for 1 h at 37°C. Huh-7 cells (Japanese Collection of Research Bioresources, 0403) were seeded to the plate and cultured for 20-28 h in 5% CO_2_ at 37°C. After 1-day culture, the cells were lysed and treated by luciferase substrate (Perkinelmer, 6066769). The chemiluminescence signals were recorded by PerkinElmer Ensight (PerkinElmer, HH3400). All neutralization assays were conducted in at least two independent experiments. Each batch of neutralization assay consists of 9 monoclonal neutralizing antibodies with known epitopes and activity to serve as controls for standardization purposes.

#### Surface plasmon resonance

SPR experiments were performed on the Biacore 8K (GE Healthcare) platform, as described previously ^9^. Human ACE2 was immobilized on a CM5 sensor chip and purified RBDs of SARS-CoV-2 variants (BA.2.75, BQ.1.1, XBB/XBB.1, and XBB.1.5) were injected. The response units were recorded by Biacore 8K Evaluation Software (GE Healthcare) at room temperature, and the raw data curves were fitted to a 1:1 binding model using Biacore 8K Evaluation Software (GE Healthcare).

**Figure S1.**
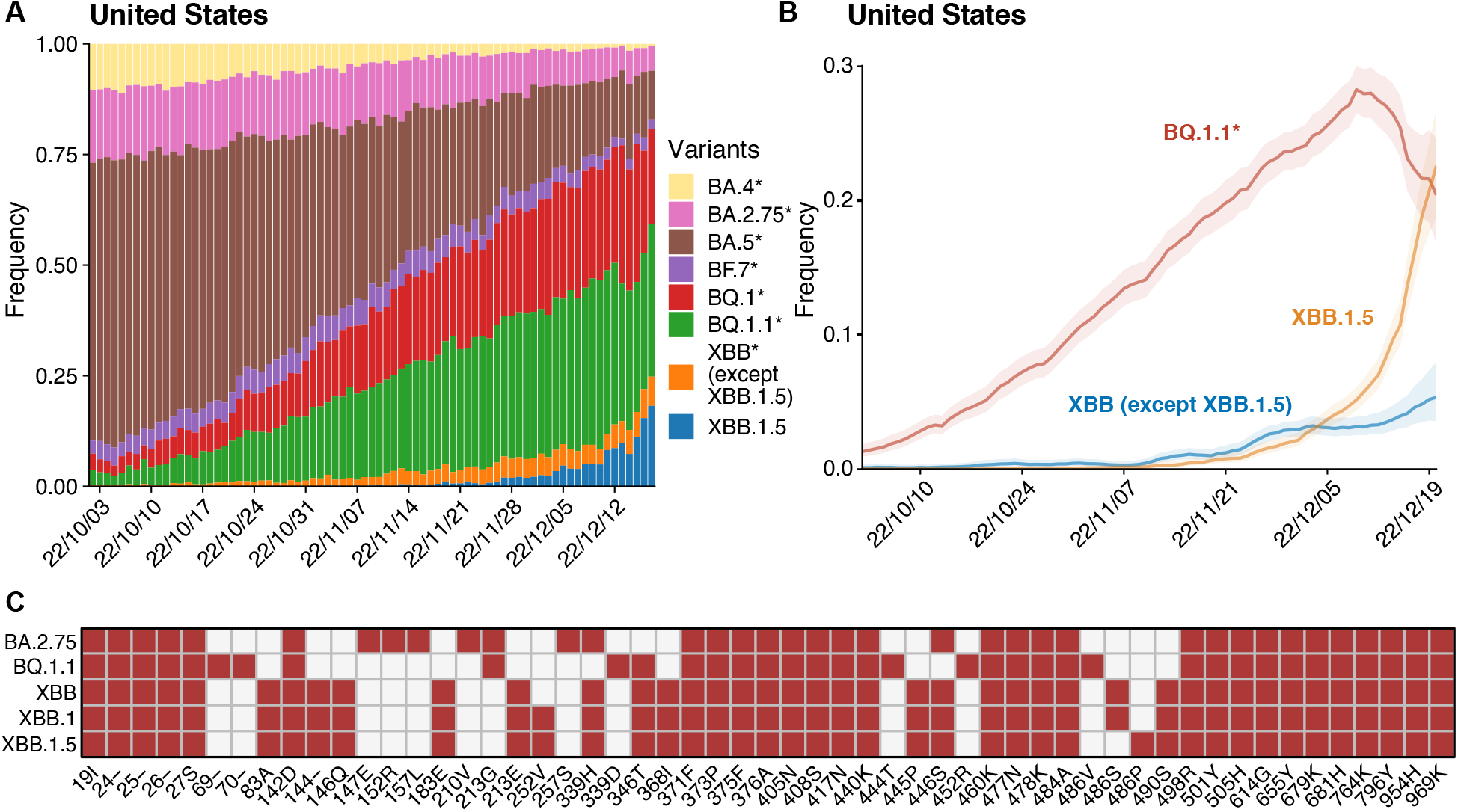
XBB.1.5 exhibited growth advantage compared to other evasive strains. (A) Dynamics of the proportion of major SARS-CoV-2 variants circulating in the United States in the past 3 months. Lightweight API for Sequences (LAPIS) is used to query the frequencies of strains. (B) XBB.1.5 showed a growth advantage compared to BQ.1.1 lineage and other XBB sublineages. Data were downloaded from CoV-Spectrum (https://cov-spectrum.org). (C) Spike mutations carried by variants involved in this study. The corresponding mutations are colored red.

**Figure S2.**
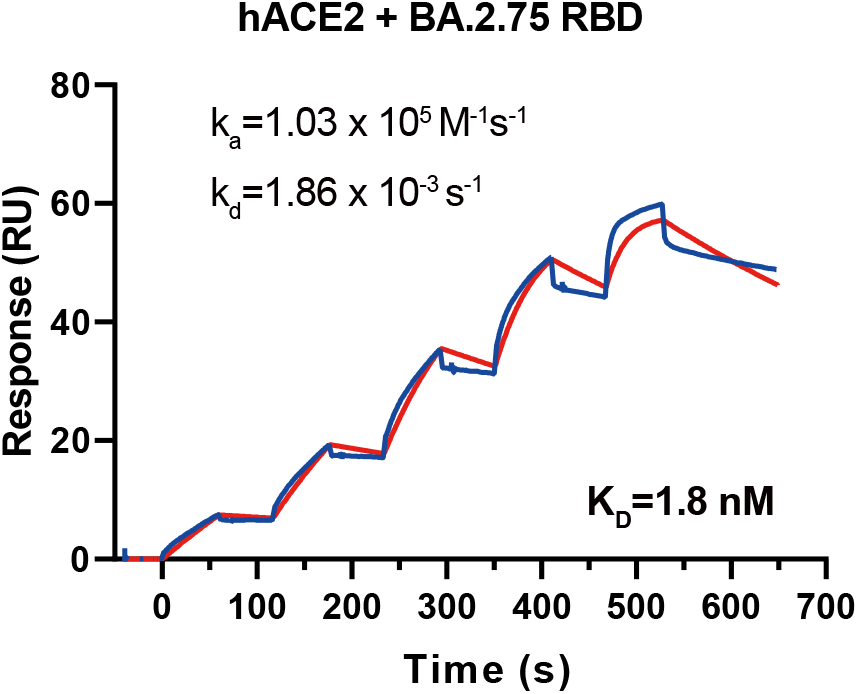
SPR sensorgram measuring the affinity of BA.2.75 RBD to human ACE2.

**Table S1 Information of the SARS-CoV-2 BTI convalescent individuals involved in this study.**

## Notes

### Summary of Updates

Additional method descriptions added

